# Identification of candidate flowering and sex genes in white Guinea yam (*D. rotundata Poir*.) by SuperSAGE transcriptome profiling

**DOI:** 10.1101/626200

**Authors:** Gezahegn Girma, Satoshi Natsume, Anna Vittoria Carluccio, Hiroki Takagi, Hideo Matsumura, Aiko Uemura, Satoru Muranaka, Hiroko Takagi, Livia Stavolone, Melaku Gedil, Charles Spillane, Ryohei Terauchi, Muluneh Tamiru

**Author notes:** Corresponding Author Email addresses (SM). Current address: Istituto per la Protezione Sostenibile delle Piante, Consiglio Nazionale delle ricerche, Bari, Italy (Gezahegn Girma). Current address: Department of Botany and Plant Pathology, Purdue University, West Lafayette, Indiana, USA. Current address: Centre for AgriBioscience, Department of Animal, Plant and Soil Science, La Trobe University, Melbourne, Bundoora, Australia (Muluneh Tamiru).

## Abstract

Dioecy (distinct male and female individuals) combined with scarce to non-flowering are common features of cultivated yam (*Dioscorea* spp.). However, the molecular mechanisms underlying flowering and sex determination in *Dioscorea* are unknown. We conducted SuperSAGE transcriptome profiling of male, female and monoecious individuals to identify flowering and sex-related genes in white Guinea yam (*D. rotundata*). SuperSAGE analysis generated a total of 20,236 unique tags, of which 13,901 were represented by a minimum of 10 tags. Of these, 88 tags were significantly differentially expressed in male, female and monoecious plants. Of the 88 differentially expressed SuperSAGE tags, 18 corresponded to genes previously implicated in flower development and sex determination in multiple plant species. We validated the SuperSAGE data with quantitative real-time PCR (qRT-PCR)-based analysis of the expression of four candidate genes. Our findings suggest that mechanisms of flowering and sex determination are likely conserved in *Dioscorea*. We further investigated the flowering patterns of 1938 *D. rotundata* accessions representing diverse geographical origins over two years, revealing that over 85% of the accessions are either male or non-flowering, and that less than 15% are female, while monoecious plants are rare. Intensity of flowering appeared to be a function of sex, with male plants flowering more abundantly than female ones. Candidate genes identified in this study can be targeted with the aim to induce regular flowering in poor to non-flowering cultivars. Findings of the study provide important inputs for further studies aiming to overcome the challenge of flowering in yams and to improve the efficiency of yam breeding.

## Introduction

White Guinea yam, *Dioscorea rotundata* Poir., is the most preferred and widely cultivated yam species in West-Africa [1]. Despite its considerable economic and socio-cultural importance, genetic improvement of cultivated yam remains difficult and slow due to its dioecy and poor to non-flowering nature [2]. Dioecy, the presence of distinct male and female individuals, is one of the major characteristics of the genus *Dioscorea* [3]. A major breeding challenge associated with dioecy is synchronizing flowering time for difficult when making genetic crosses. In addition, many cultivars of *D. rotundata* rarely flower, while a significant proportion of those that do flower seldom set fertile seeds [4].

Mechanisms of sex determination in plants have been investigated in over 40 angiosperm species that identified of heteromorphic and monomorphic sex chromosomes as well as XY and ZW sex-determination systems [5,6]. Due to the large number and small sizes of chromosomes, the identification of sex chromosomes at cytological level is difficult in *Dioscorea* [8, 9]. In *D. tokoro,* a diploid wild species, a heterogametic (XY: male) and homogametic (XX: female) sex has been proposed based on segregation patterns of AFLP markers tightly linked to sex [5,6]. However, a more recent study has revealed that sex determination in the cultivated African species of *D. rotundata* follows the ZZ (male) and ZW (female) system [7]. In the same study, a candidate chromosomal region associated with sex was determined and a diagnostic marker for sex determination developed. However, gene(s) functionally responsible for sex determination are yet to be identified in yam species.

In angiosperms, flower development involves gene expression and pathway changes in vegetative meristems, leading to conversion to flowering meristems in response to environmental cues and developmental signals [8]. Molecular genetic analyses in eudicots such as *Arabidopsis thaliana* and *Antirrhinum majus* have led to the identification of several floral organ-identity genes, which were originally grouped into three classes (class A, B, and C) based on the floral organ identity they specify [9,10]. The class A, B, and C genes are required for the development of sepals and petals, petals and stamens, and stamens and carpels, respectively. Class A and C genes are mutually antagonistic. As master regulators of floral organ identity, plant MADS-box transcription factors are central to the ABC model [11]. Further studies revealed that additional gene classes, class D and E, are important for ovule and organ development, respectively, leading to a modified ABCDE model for flower development [12].

A range of factors can regulate sex differentiation in angiosperms including genetic [13], epigenetic and environmental [14], as well as physiological regulation by phytohormones [15]. Understanding the molecular and genetic mechanisms of flowering is essential for efficient plant breeding. Recently, whole genome sequencing of *D. rotundata* has allowed the identification of a genomic region associated with sex [7]. This finding is expected to open opportunities for dissecting the molecular mechanisms regulating flowering and sex determination in yam. Identification of flowering and sex-related genes is critical to scaling up utilization of the available yam germplasm in breeding programs.

Various sequencing-based transcriptome profiling techniques are available for gene expression profiling, novel gene discovery, and genome annotation studies. These include Serial Analysis of Gene Expression (SAGE), which is based on unique short sequence tags 14–15 bp [16], LongSAGE which uses a different type IIS enzyme, *Mme*I, to generate 21-bp fragments from each transcript [17], Robust-LongSAGE (RL-SAGE) [18], Expressed Sequence Tag Analysis (EST) Analysis (Nielsen 2006) [19], Digital Gene Expression TAG (DGE-TAG), DeepSAGE [19], and RNA-Seq [20]. The next generation sequencing (NGS)-based and high-throughput SuperSAGE for tag-based gene expression profiling involves sequencing of longer fragments (26-bp) and simultaneous analysis of multiple samples by using indexing (barcoding) [21]. The longer tags generated by SuperSAGE compared to the relatively shorter tag reads obtained by other NGS-based techniques such as DGE-TAG significantly improve the accuracy of tag-to-gene annotation [21].

In this study, we applied SuperSAGE to analyze for transcriptome changes in flowers at early stage of development from *D. rotundata* accessions representing three flowering groups (male, female and monoecious) and identified differentially expressed genes. We show that majority of these genes correspond to known genes, including genes that regulate flowering and sex determination in multiple species. Our findings suggest that known mechanisms underlying flowering and sex determination are likely to be conserved in yams.

## Materials and Methods

### Morphological characterization

A total of 1938 accessions of *D. rotundata* planted for routine field maintenance under IITA Genetic Resources Center were characterized based on 12 morphological traits as well as their flowering pattern recorded for two consecutive seasons in 2010 and 2011 seasons. The 12 morphological traits were selected among the yam descriptors jointly developed by the former International Plant Genetic Resources Institute (current known by its new designation Bioversity International) and the International Institute of Tropical Agriculture [22]. Detailed description of the traits used is provided in S1 Table.

### Sampling and RNA extraction

Tubers of seven accessions including two male, two female, and three monoecious (Table 1) selected based on their consistency of flowering over two years were planted in pots in a screen house. Samples for RNA extraction were collected from early stage flowers (Fig 1). Total RNA was extracted using the Qiagen RNeasy plant mini kit according to manufacturer’s protocol (Qiagen, Venlo, the Netherlands). On-column DNAase treatment was performed to remove contaminating DNA.

**Table 1.** Summary of SuperSAGE tags generated by Illumina sequencing of *D. rotundata* accessions representing different flowering groups

| Accession | Sex | Total tags | Unique tags | Non-singleton tags |
| --- | --- | --- | --- | --- |
| TDr3631 | Male | 1,251,361 | 17,773 | 16,602 |
| TDr2965 | Male | 1,460,689 | 18,234 | 17,408 |
| TDr4087 | Female | 1,049,552 | 18,620 | 17,853 |
| TDr1679 | Female | 560,257 | 17,534 | 15,887 |
| TDr4162 | Monoecious | 1,209,229 | 18,032 | 17,146 |
| TDr1506 | Monoecious | 1,309,189 | 18,746 | 18,177 |
| TDr1819 | Monoecious | 1,492,246 | 18,918 | 18,403 |

**Figure 1.**
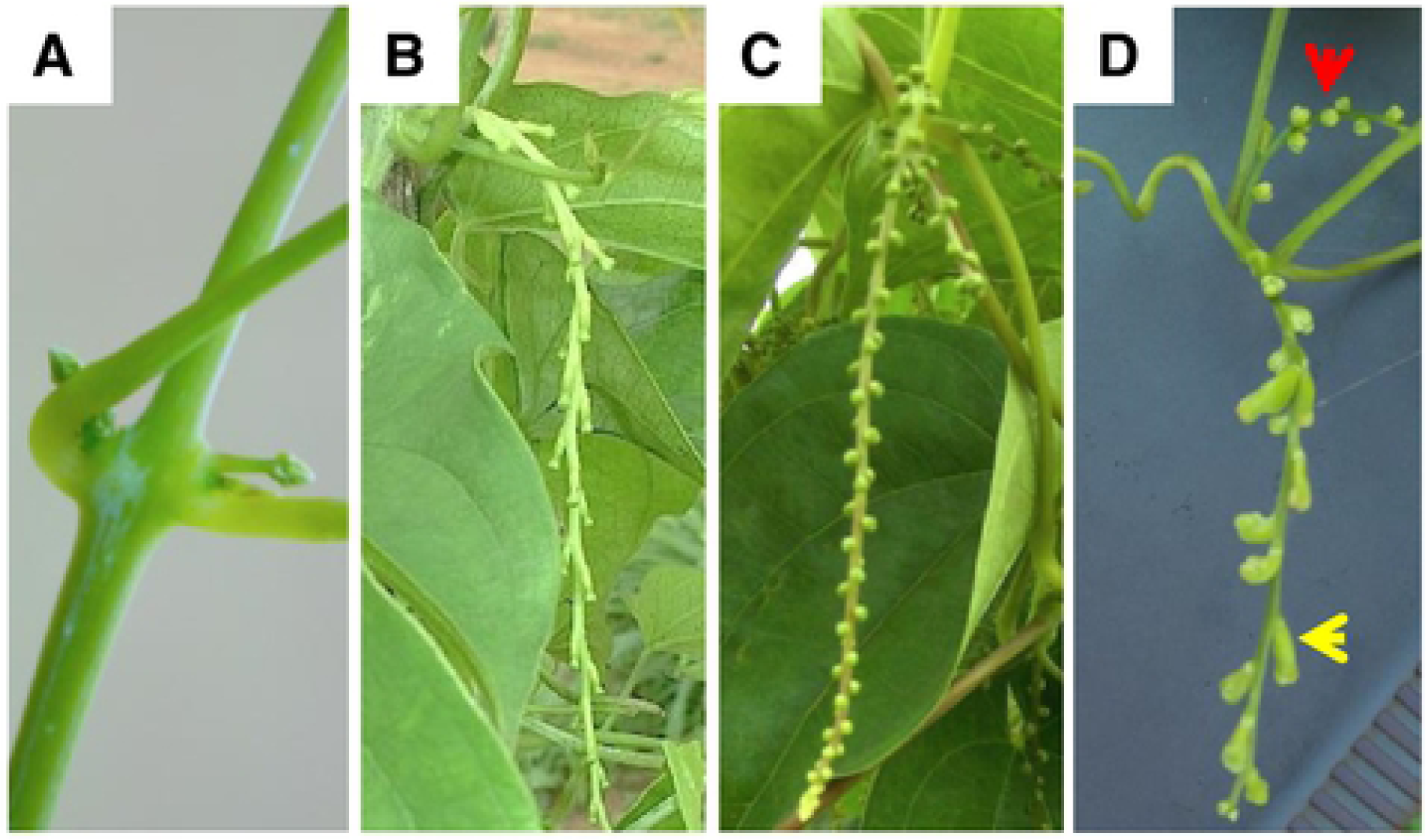
Flower types in yam (D. *rotundata*). (A) Example of flowers at early growth stage which also corresponded to the stage at which samples were collected for SuperSAGE analysis, (B) female flowers, (C) male flowers, and (D) monoecious plant with separate male (red arrow) and female (yellow arrow) flowers.

### SuperSAGE: library preparation and sequencing

SuperScript II double-strand cDNA synthesis kit was employed for cDNA synthesis using the biotinylated adapter-oligo (dT) primer (5’-bio-CTGATCTAGAGGTACCGGATCC-CAGCAGTTTTTTTTTTTTTTTTT-3’).

Synthesized cDNA was purified using Qiagen PCR purification Kit. The library was prepared following the protocols of Matsumura et al. [21]. Briefly, the NlaIII anchoring enzyme was used for sample digestion, and the fragments (NlaIII-digested cDNA) were bound to streptavidin-coated magnetic beads (Dynabeads streptavidin M-270). Non-biotinylated cDNA fragments were removed by washing, and then adapter 2 was ligated to digested cDNA fragments bound to the magnetic beads. The type III restriction enzyme EcoP151 was used for digestion of adapter 2-cDNA after washing. Adapter2-26bp fragments were further ligated to adapter 1 (that are specific for each of the samples). The adapter prepared following procedure described by [21] were used.

The adapter2-tag-adapter1 ligates was amplified using Phusion High polymerase and GEX primers (5’-AATGATACGGCGACCACCGACAGGTTCAGAGTTCTACAGTCCGA-3’ and 5’-CAAGCAGAAGACGGCATACGATCT-3’). The amplification program was 98°C for 1min, 10 cycles at 98°C for 35 sec, and 60°C for 30 sec. The PCR product consisting eight tubes per sample was pooled and concentrated using Qiagen MinElute reaction purification kit. The amplification product was run on an 8% non-denaturing polyacrylamide gel. After staining with SYBR green (Takkara Bio), the band around 123-125-bp size was cut out from the gel, and DNA purified after its elution from the gel pieces. The purified PCR product from each sample was analyzed for its quantity and quality on an Agilent Bioanalyzer 2100.

As a next step the PCR product was cloned using invitrogen: - zeroblunt Topo PCR cloning kit for sequencing and later transformation using one shot chemical transformation protocol. Colony PCR was done by using 2x colony PCR mixture and purified using QIAGEN PCR purification kit. Purified and mixed PCR products were applied to Illumina Genome Analyzer II for sequencing reactions following the manufacturers protocol.

### Real-time quantitative reverse transcription PCR (qRT-PCR)

Total RNA was extracted from female and male flowers primordia using the CTAB extraction protocol as described by (Dellaporta 1983) [23] with some modifications. Approximately 500 mg tissues were grinded using mortar and pestle in liquid nitrogen. A pre-heated 1000 uL of CTAB extraction buffer (2% CTAB, 2% PVP-40, 20 mM Tris–HCl pH 8.0, 1.4 M NaCl, 20 mM EDTA) was added to each sample and incubated at 65°C for 15 minutes, vortexed every 5-minutes and centrifuged at 15000 rpm at 4°C for 5 minutes. The aqueous top layer was transferred and purified with one volume of chloroform: isoamyl alcohol (24:1) and centrifuged for 10 minutes at 15000 rpm. The supernatant was mixed with 0.6 volume of cold isopropanol to precipitate the nucleic acid, mixed gently by inversion and centrifuged at 15000 rpm for 20 minutes. The precipitated pellet was washed with 100 uL of cold 70% ethanol and centrifuged for 5 minutes and air dried. RNA preparations were subjected to on-column DNase digestion (RNA clean and concentrator kit; ZymoResearch) and reverse-transcribed in the presence of random hexamers (LunaScript®RT SuperMix Kit, NEB *Inc*.) in 20 μl of total reaction volume containing 1 μg of total RNA and incubated at 55°C for 15 minutes following manufacturer’s instructions. Primer pairs used for qRT-PCR were designed for the four selected candidate genes (DnaJ-like protein, TK, GST and PIF3) with the support of the Integrated DNA Technologies’ PrimerQuest software and are listed in S2 Table. Three different housekeeping genes; Adenine phosphoribosyl transferase (APT), Beta-Tubulin (Tub) and TIP41-like family protein (TIP41), were tested. Based on the results of melt curve and standard curves, Beta-Tubulin was chosen as the reference endogenous gene for all targets quantification. RT-PCR reaction volumes were set up to 12.5 μL containing Luna Universal qPCR Master Mix (New England Biolabs *Inc*.), 2 μl of a 1:10 dilution of cDNA reaction, and 400 nM each of corresponding forward and reverse primers. Three different biological replicates represented by total RNA extracted from three individual plants and reverse transcribed into cDNA by LunaScript®RT SuperMix Kit (New England Biolabs *Inc*.) were used for statistical analysis of the quantification. Each cDNA sample was amplified in duplicate on a single 96-well optical plate using LightCycler480 (Roche). The cycling profile consisted of 95°C for 10 min followed by 40 cycles of 15 s at 95°C and 60 s at 60°C, as recommended by the manufacturer. Immediately after the final PCR cycle, a melting curve analysis was done to determine specificity of the reaction.

## Data analysis

For analysis of data on phenotypic traits, multiple correspondence analyses (MCA) was performed using FactoMineR package [24] in R software [25] to detect the underlying structure of morphological variables and its correlation with flowering patterns in the data set.

For SuperSAGE data analysis, sequence reads obtained from the different libraries were first sorted into different flowering groups based on their respective index sequences. Then, the subsequent extraction of SuperSAGE tags from reads was conducted using a script written in Perl. The R package edgeR [26] was used to determine differentially expressed tags across different pairs of flowering patterns (male vs female, male vs monoecious and female vs monoecious). Tags with very low counts were filtered out and tags that were expressed in at least one sample from each flowering group were considered for the analysis. Additionally, since two replicate samples were sequenced from each flowering group, tags that expressed in both replicates were considered. The expression cutoff was 100 counts per million (CPM), corresponding to a read count of about 5 for each tag.

For annotation of SuperSAGE tags, the selected tags were first aligned to the draft *D. rotundata* scaffold sequence, followed by extraction of 2000-bp upstream sequences, which were finally used as queries for BLAST search against the non-redundant (nr) database of the National Center for Biotechnology Information (NCBI) and the Universal Protein Resource (UniProt).

For qRT-PCR analysis, quantities of RNA accumulation levels were calculated as relative quantification (RQ) values using the comparative cycle threshold (Ct) (2– ΔΔCt) method [27]. Before quantitative analyses, validation experiments were carried out to confirm equal amplification efficiencies between reference and target genes and the applicability of the comparative method. Assessing the relative amplification efficiencies was achieved by running standard curves for each amplicon (i.e., five serial log10 dilutions of starting cDNA were amplified and Ct values of target and reference genes were measured in triplicate and plotted against the log of the input cDNA amount). The efficiencies were considered comparable when falling within a range of 100 ± 10%, corresponding to a curve slope of –3.3 ± 0.33 [28], and all primers pairs that did not allow PCR performances matching these criteria were discarded.

## Results

### Variation in flowering patterns and morphological traits in *D. rotundata* genebank accessions

The flowering patterns of 1938 *D. rotundata* accessions collected primarily from the main yam growing regions of West and Central Africa were assessed under field conditions at IITA over two consecutive growing seasons in 2010 and 2011. *D. rotundata* accessions are easily distinguished based on their flowers as female, male, or monoecious (unisexual female and male flowers on the same plant) (Fig 1). In 2010, majority of the accessions (996, 51.4%) failed to flower, while 745 (38.4%), 170 (8.8%), and 27 (1.4%) were classified as male, female, and monoecious, respectively (Fig 2). In the 2011 growing season, 939 (48.5%) accessions were male, followed by 630 (32.5%) non-flowering, 287 (14.8%) female, and 82 (4.2%) monoecious accessions (Fig 2b). Most accessions were consistent over the two seasons with respect to flowering, while some were not. About 326 (16.8%) male, 169 (8.7%) female and 43 (2.2%) monoecious accessions failed to flower at least in one of the seasons, hence the discrepancy observed in the proportion of accessions with different sexes between the two seasons (Fig 2). Overall, majority of *D. rotundata* accessions maintained at IITA were either male or non-flowering, while the female accessions represented less than 15% of the collection. The results indicate that monoecious types are very rare in *D. rotundata*.

**Figure 2.**
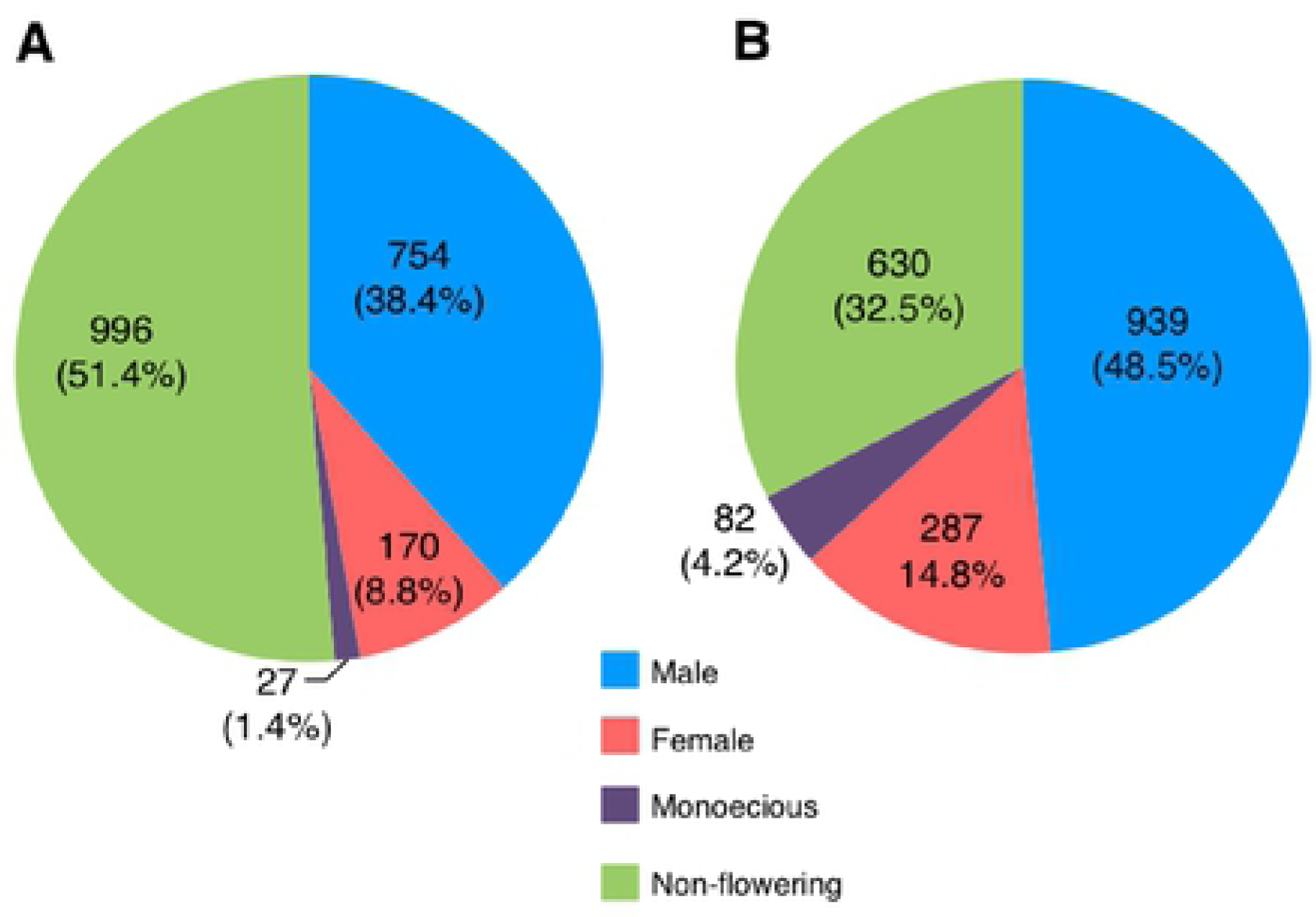
Sex distribution in yam (D *rotundata)* accessions. The proportion of male, female, monoecious, and non-flowering accessions among 1938 genebank accessions in (A) 2010 and (B) 2011 growing seasons.

In addition to flowering characteristics, we collected categorical data on 14 selected morphological traits over the same period (S1 Table). The data was subjected to multiple correspondence analysis (MCA), generating three major clusters that mainly reflected sex of the accessions, suggesting an overall correlation between sex and the selected morphological traits (Fig 3). The first cluster included a distinct group consisting of non-flowering accessions that were distinguished by traits such as purplish green stem, non-waxy stem, stem with non-barky patches, dark green leaf color, and hastate leaf shape. A second cluster composed entirely of male accessions was correlated with purplish green stem, presence of barky patches, non-waxy stem, dark green leaf, and sagittate leaf shape. The third group was composed of a mixture of male, female and monoecious flowering accessions that were identified by waxiness, absence of barky patches, either green, brownish green or purple stem color, and pale green or green leaf color as distinct traits.

**Figure 3.**
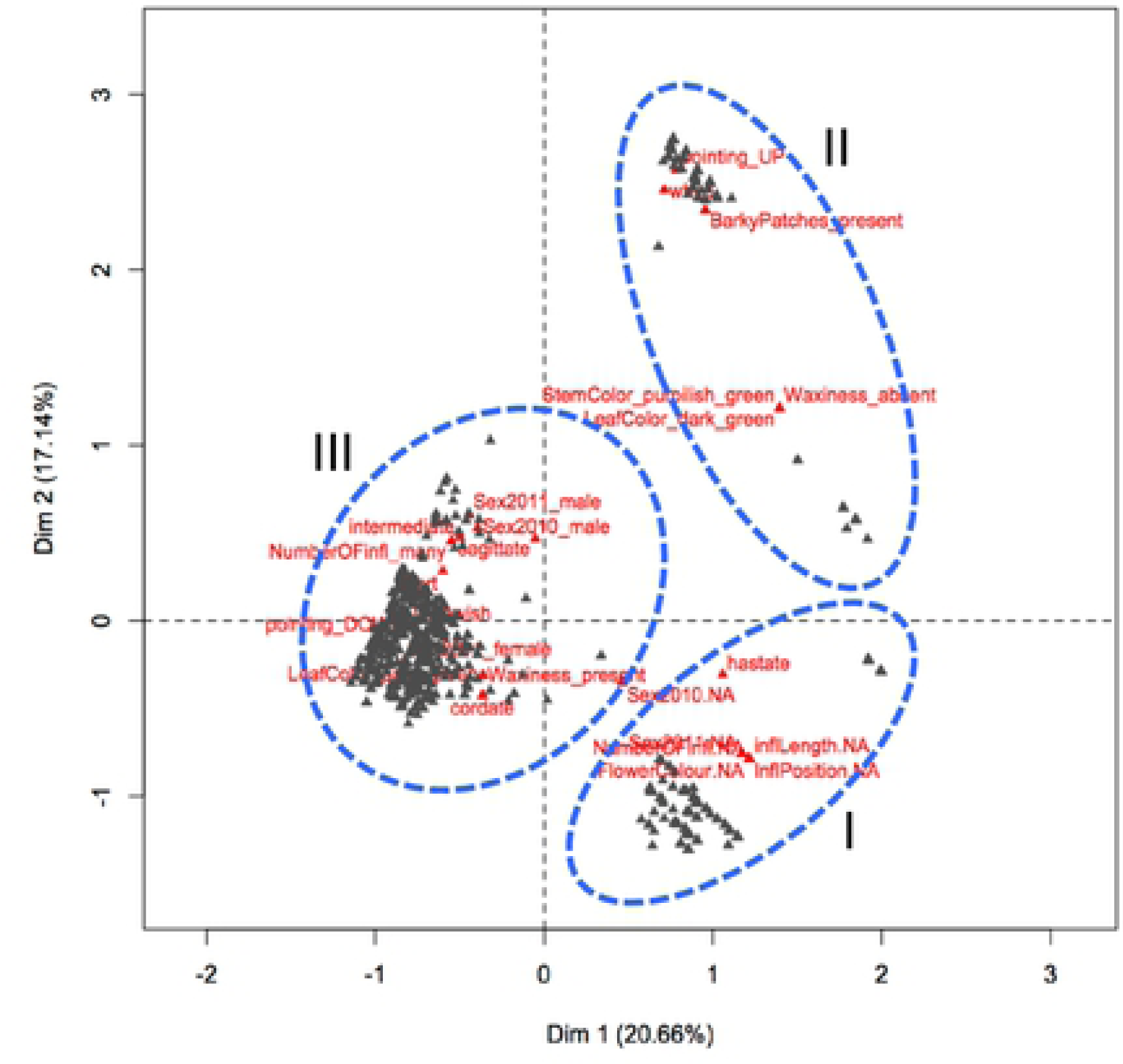
Multiple correspondence analysis (MCA) of sex type and phenotypic traits in yam (D. *rotundata).* The pattern of relationship between individual plants (black triangles) and the 20 most discriminant morphological traits (red triangles) are provided. The blue circles with broken lines represent the three main cluster: Cluster I = non-flowering accessions; Cluster II = male accessions; Cluster Ill = male, female, and monoecious accessions.

### Generation and analysis of SuperSAGE tags

SuperSAGE libraries representing early stage floral samples of male, female, and monoecious flower groups were multiplexed and sequenced on a single lane of an Illumina Genome Analyzer IIx, generating a total of 8,332,523 tags after quality control (QC) based on sequence read length, adapter dimmer, and sequencing error rates (Table 1). Following sorting of tags into the different flowering groups based using Adapter1 sequences and removal of singleton tags, 20,236 unique non-singleton tags were obtained. Of these, 6,335 tags that were ten or less in number in each library were removed, while the remaining 13,901 abundant tags were retained for further analysis (S3 Table). Of these, 5985 (43%) tags were shared among the three flowering groups, whereas other tags were specific to male (1855), female (1648) or monoecious (765) flowers (Fig 4). The remaining 2650 (19.0%), 378 (2.7%), and 620 (4.5%) were shared between male and female, male and monoecious, and female and monoecious groups, respectively (Fig 4).

**Figure 4.**
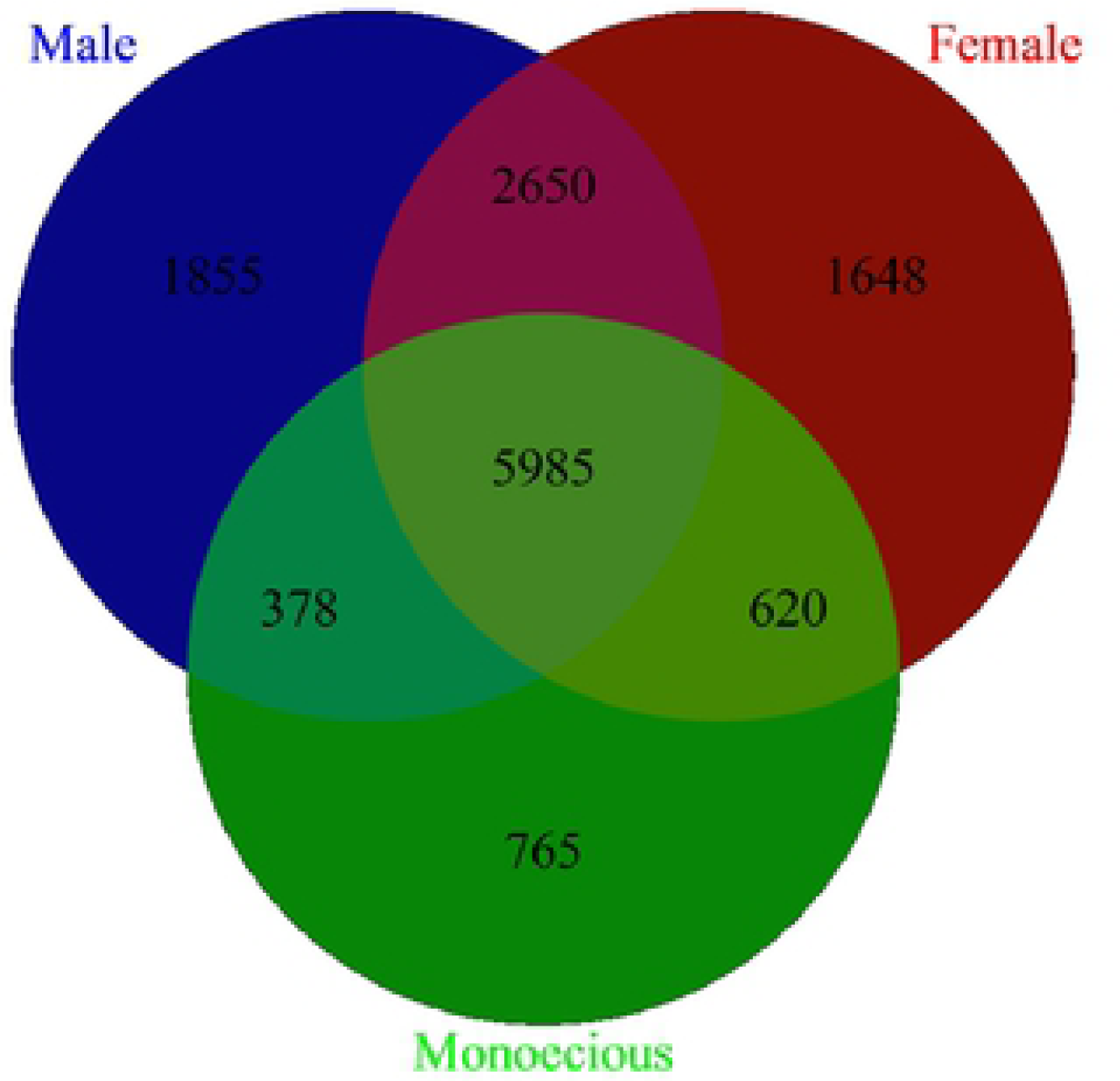
Distribution of SuperSAGE tags in flower buds of three yam (*D. rotundata*) sex types. The unique tags, as well as tags shared among male, female and monoecious p ants were presented.

### Identification of differentially expressed genes across different flowering groups of yam

The fold change of differential expression, and gene abundance (count per million) of 13,901 unique tags was compared across the different flowering groups. A total of 100 tags were differentially expressed with *p* and FDR values less than 0.01 (Fig 5; S4 Table). The number of differentially expressed SuperSAGE tags between male vs. female, male vs. monoecious and female vs. monoecious groups were 13, 67 and 20, respectively (S4 Table). Of the 13 tags differentially expressed between male and female groups, five were highly expressed in male while the remaining eight were expressed in the female sex type. Similarly, the male vs monoecious group comparison revealed that 25 tags were highly represented in male, and 42 were abundant in monoecious sex type. For the female vs monoecious flower group, 11 and nine tags were expressed highly in female and monoecious flowers, respectively. The tag abundance estimated by the average logCPM (counts per million) ranged from a minimum of 6.36 to 11.88 for all the differentially expressed tags (Fig 5 and S4 Table).

**Figure 5.**
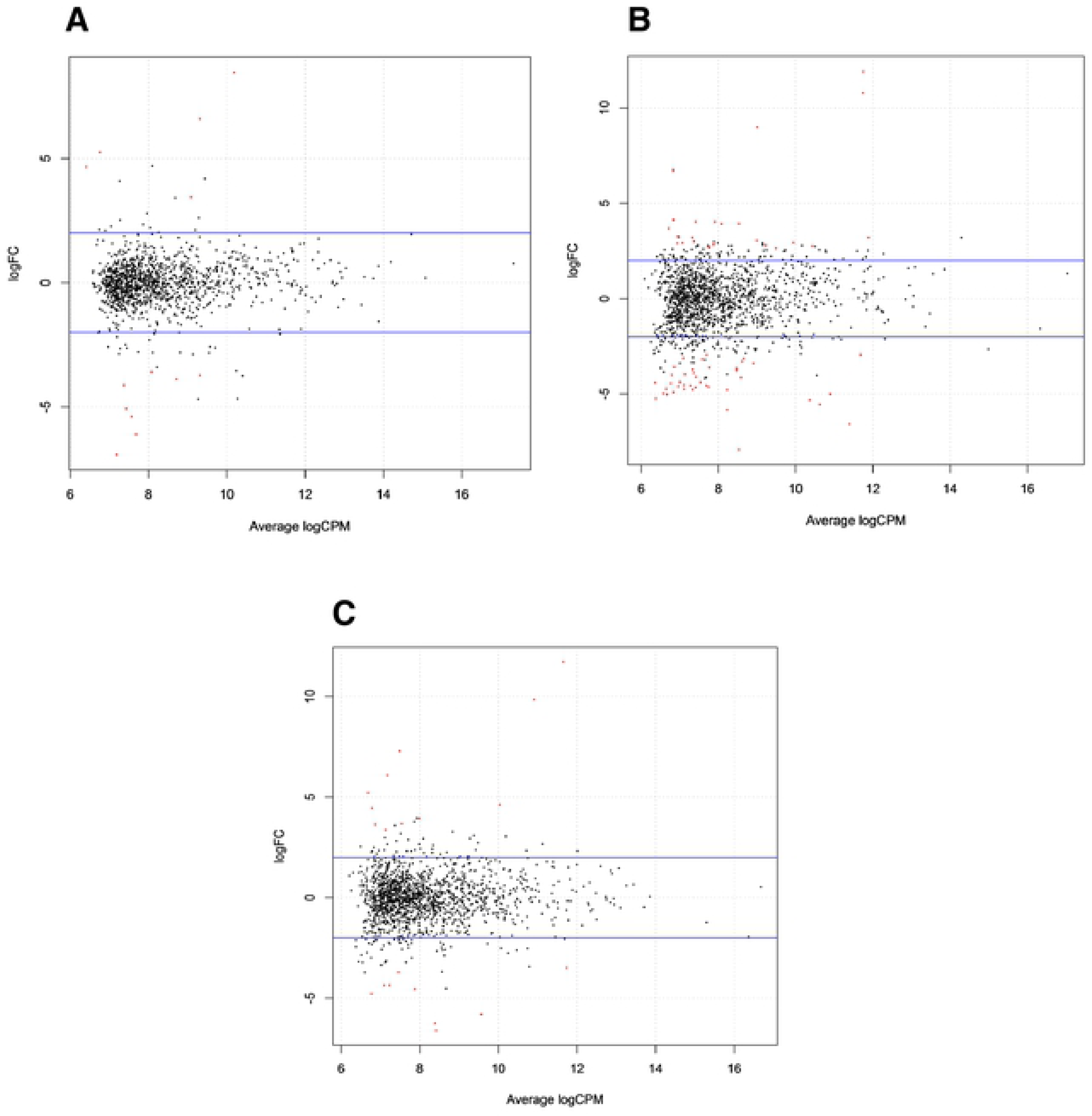
Differentially expressed genes in flower buds of three yam (D. *rotundata)* sex types. The abundance of genes differential expressed in (A) male and female, (B) male and monoecious, and (C) female and monoecious flowers buds are shown. Differentially expressed tags are represented by red dots. Fold change values between groups are plotted against average log expression values (standardized read counts) The logFC indicates the fold changes of different al expression whereas logCPM indicate count per million or tag/gene abundance The horizontal blue lines represent 4 fold changes.

### Annotation of SuperSAGE tags

Of the 100 differentially expressed tags, 88 tags were unique. These unique tags were aligned to *D. rotundata* draft scaffold sequence to extract 2000-bp upstream sequences for use as a query for tag annotation by BLAST search against the National Center for Biotechnology Information (NCBI) and Universal Protein Resource (UniProt) databases. Of the 88-unique sequences, 87 could be aligned to the draft *D. rotundata* sequence, and of these 72 matched sequences available in the NCBI and UniProt databases. Sequences obtained for fifteen (17.24%) tags did not match sequences available in the two databases, while eight (9.09%) tags generated high e-values. Of the 72 tags that could be annotated, 14 (19.5%) corresponded to proteins of unknown function, unnamed proteins, uncharacterized proteins or hypothetical proteins (S5 Table). However, a set of 18 (25.0%) tags representing 16 unique genes corresponded to genes that have been previously described as having a role in flower development and/or being expressed in flowers in multiple species (Table 2).

**Table 2.**
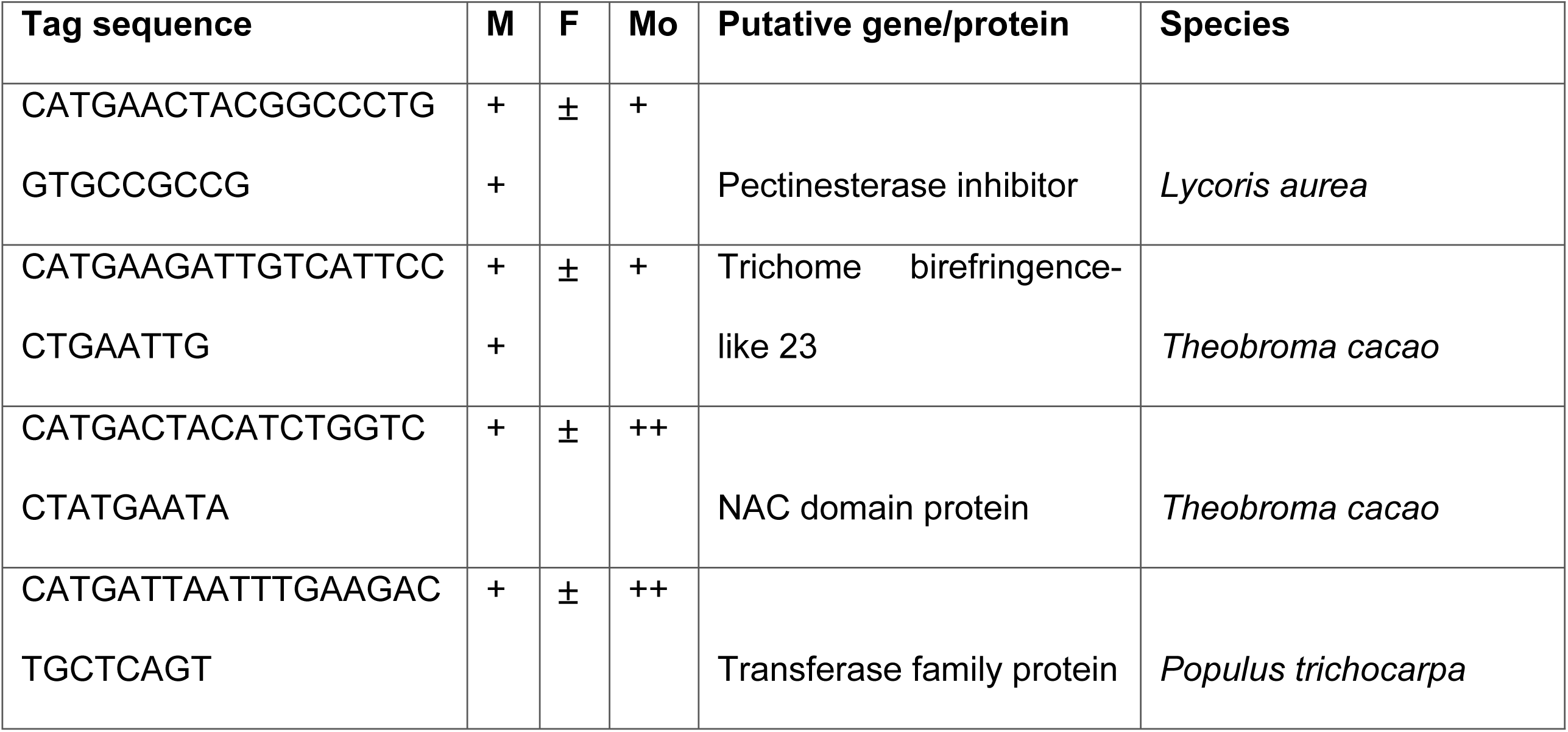
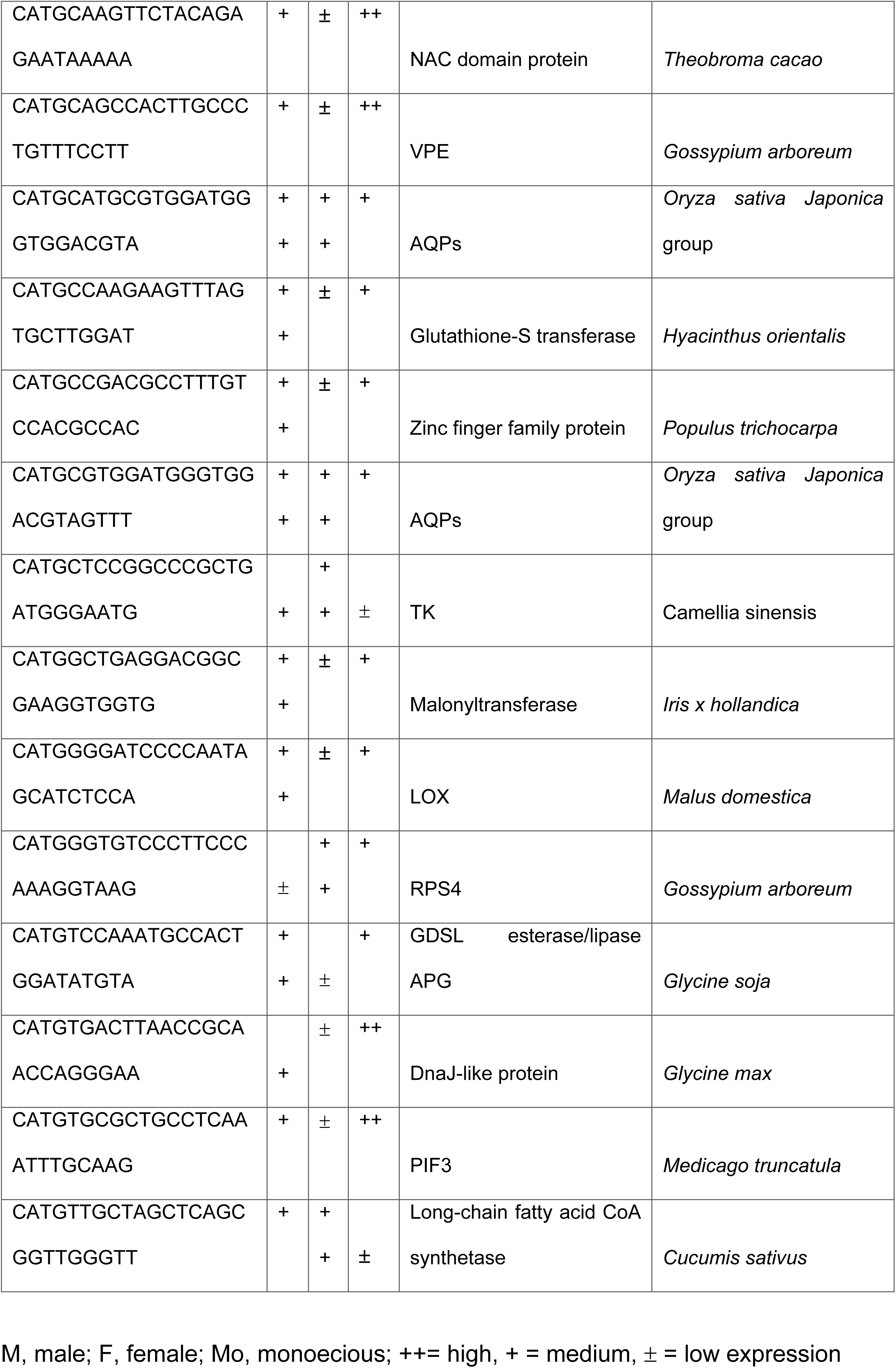
Selected SuperSAGE tags generated from *D. rotundata* and corresponding to genes expressed in flowers or involved in flower development in multiple species

Among 16 genes reported to have involvement in flower and flower development, four and twelve had high and low expression in female, respectively. Whereas eight, seven and one in male and five, nine and two in monoecious flowers had high, medium and low expression, respectively (Table 2). Our results indicate that 16, 15, and four genes were preferentially expressed in male, monoecious, and female flowers, respectively. However, only one gene was differentially expressed across all the flowering groups, whereas two, three and 14 genes were differentially expressed between female and monoecious, male and female, and male and monoecious flowers, respectively.

Following blastn of the 88 deferentially expressed tags to *D. rotundata* scaffolds, a single hit to eight was found across 70 tags. The remaining 18 tags had no hit (S6 Table). Further conversion of the scaffold position of the tags to recently published [7] pseudo chromosome as “TDr96_F1_Pseudo_Chromosome_v1.0.fasta” detected distribution of the tags across all chromosomes except in chromosome_01. The tag count ranged from two in seven of the pseudo chromosomes to the largest tag count of 11 in chromosome_05 (S6 Table). Moreover, the gene ID and annotations of the tags were also confirmed using the published [7] gene model, “TDr96_F1_v1.0. gff3”.

### Validation of selected candidate genes by qRT-PCR

To independently validate our SuperSAGE results using a different technique, we selected four representative genes associated with flowering and included DnaJ-like protein, Transketolase (TK), Glutathione-S Transferase (GST) and Phytochrome Interacting Factor-3 (PIF3). Three different housekeeping genes previously used for qRT-PCR across different organs and developmental stages of *Dioscorea opposita* [29] including Adenine Phosphoribosyl Transferase (APT), Beta-Tubulin (Tub) and TIP41-like family protein (TIP41) were tested based on standard curve and melt curve analysis.

Each target gene was amplified in two different samples; female (F) and male flowers and in three biological replicates. Beta-Tubulin was selected in the current experiments as the most suitable and stable housekeeping gene in *D. rotundata*. qRT-PCR amplification revealed high expression of DnaJ-like and TK in F than in M (Fig 6A and 6B). Whereas, both GST and PIF3 showed higher expression level in M and vice-versa in F (Fig 6C and 6D).

**Figure 6.**
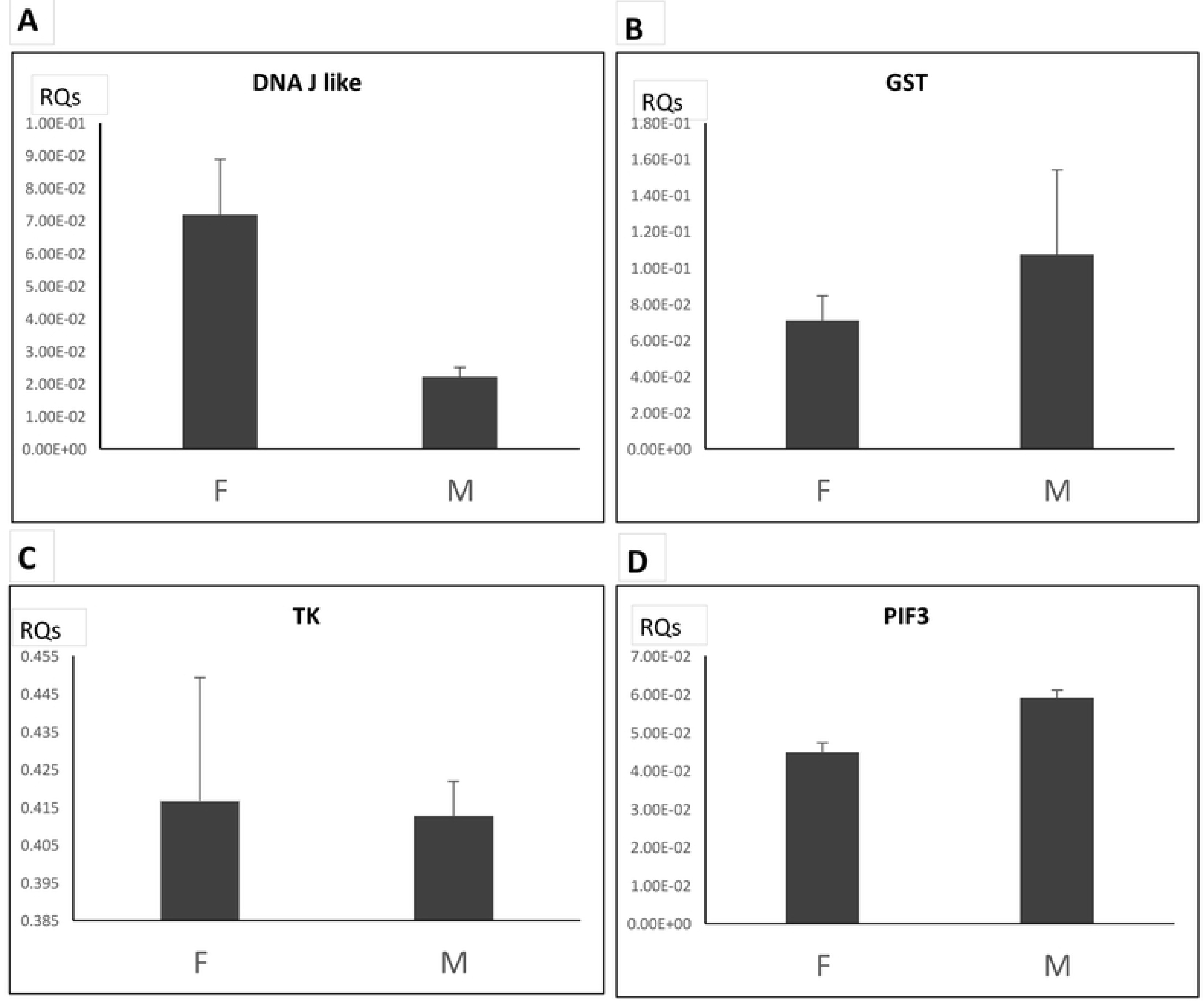
Overview of the qRT-PCR validation of selected differential expressed genes, (A) DnaJ-like protein, (B) Glutathione S-transferase-like (GST), (C) Transketolase (TK) and (D) Phytochrome interacting factor-3 (PIF3) across female (F) and ma e (M) sex types in *D rotundata.* Error bars represent the standard deviaton of ratios computed from every combination of one target with its own reference.

## Discussion

### Flowering relationships with morphological variation

A number of previous studies have shown the existence of dioecious and monoecious flowering patterns in *D. rotundata* [30-32]. Here, we have also recorded dioecious and monoecious, as well as non-flowering accessions in the *D. rotundata* germplasm maintained at IITA. The monoecious plants predominantly produced male flowers, a phenomenon previously described as trimonoecious [30]. Most of the inflorescences consisted only of male flowers, while a limited number of inflorescences contained few female flowers.

Our observations over the two seasons demonstrate that majority of the *D. rotundata* accessions being maintained at IITA are either male or non-flowering types, with a significantly lower proportion of female accessions, while monoecious accessions are rare (Fig 2). Abundance of flowering appeared to vary according to sex, as male accessions flowered profusely compared to female ones. Moreover, higher numbers of male flowers develop on monoecious plants than female flowers. A similar observation from materials collected in Benin showed that female plants are rare and produce a limited number of flowers, whereas the male flowering cultivars, whenever they flower, produce flowers in abundance [31]. Overall, flowering pattern in the two seasons were inconsistent across yam germplasm collection. This could be related to the effect of environmental factors as previously reported [30]. The variation in recorded weather data (mean monthly temperature and rainfall) in 2010 and 2011 and particularly during main flowering period (July - September) at the IITA experimental site (S7 Table) similarly suggests the effect of weather in yam flowering.

Morphological characters that could be used to distinguish the different flowering types at the earliest growth stage possible are important particularly for breeders to select germplasm and cultivars in breeding experiments. Our study revealed that a set of selected morphological traits can be used as a basis for sex prediction in *D. rotundata* accessions (Fig 3; S1 Table). The correlation between flower type, morphological traits, and ploidy level previously reported in *D. rotundata* suggested that all triploid individuals are either male or non-flowering, and display some morphological features distinct from diploid individuals [33]. This morphological traits-based prediction of sex and ploidy level provides significant practical advantages for choosing of parents for crossing in breeding trials. However, we do recognize that the correlation identified needs to be validated with repeated experiments conducted under different environmental conditions to select those traits with the highest predictive value across multiple environments.

### Identification of candidate genes associated with flowering in yams

In our study, a number of genes differentially expressed in D. rotundata flower types were identified (Table 2 and S4 Table). However, only a few of the differentially expressed genes corresponded to genes known to be associated with flowering in plants, including those that are known either for their expression in flowers, for organ or sex specific expression, for a role in the regulation of flowering time, photoperiod and the transition from vegetative to reproductive phase, or for flower color development (Table 3). Examples of candidate genes with expression in flowers include the *Pectinesterase inhibitor* and *AQPs/MIP* gene in flowers of *Arabidopsis thaliana* [34,35] the *Malonyltransferase* gene in flowers of *Salvia splendens* [36], and the *VPE* gene in *Citrus sinensis* L. during flower development [37] and expression of *Trichome birefringence-like 23* at mature pollen stage [38] and in petal, sepal, pedicel, stamen, pollen, and petal differentiation and expansion stages of *Arabidopsis thaliana* [39,40].

**Table 3.**
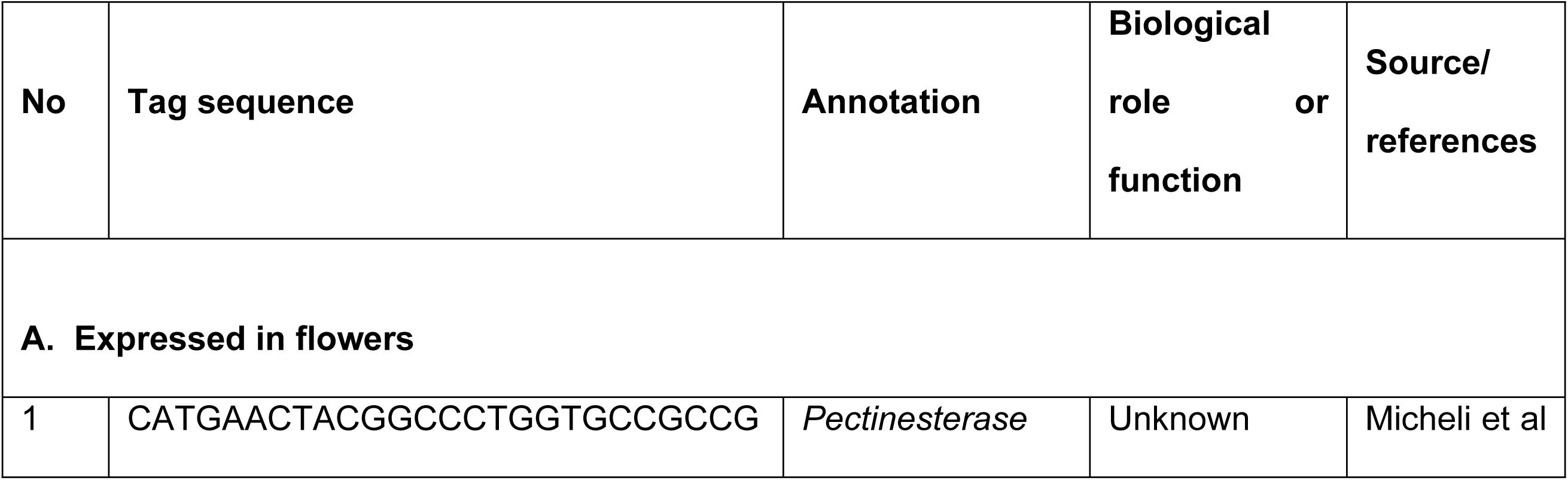
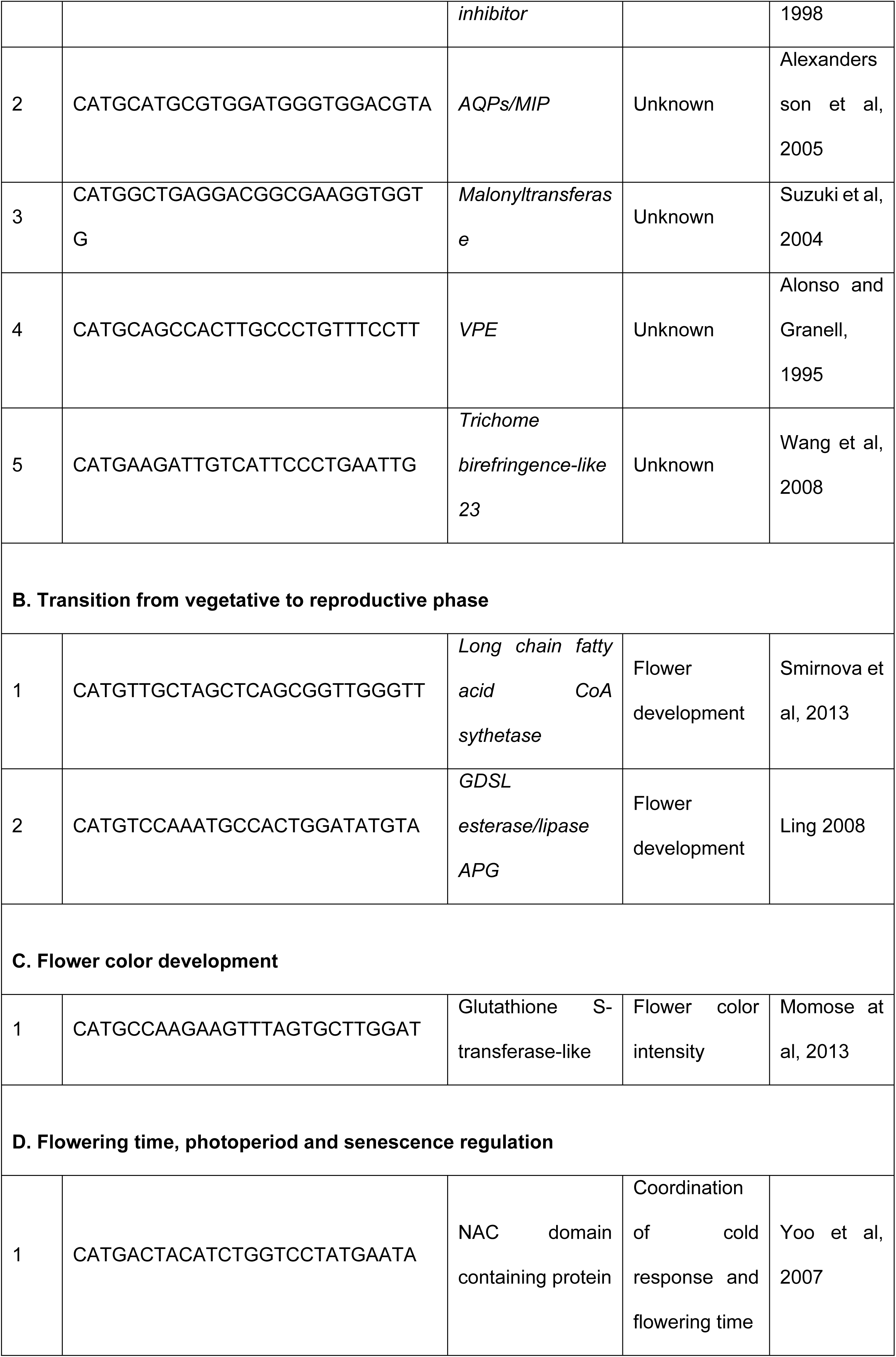
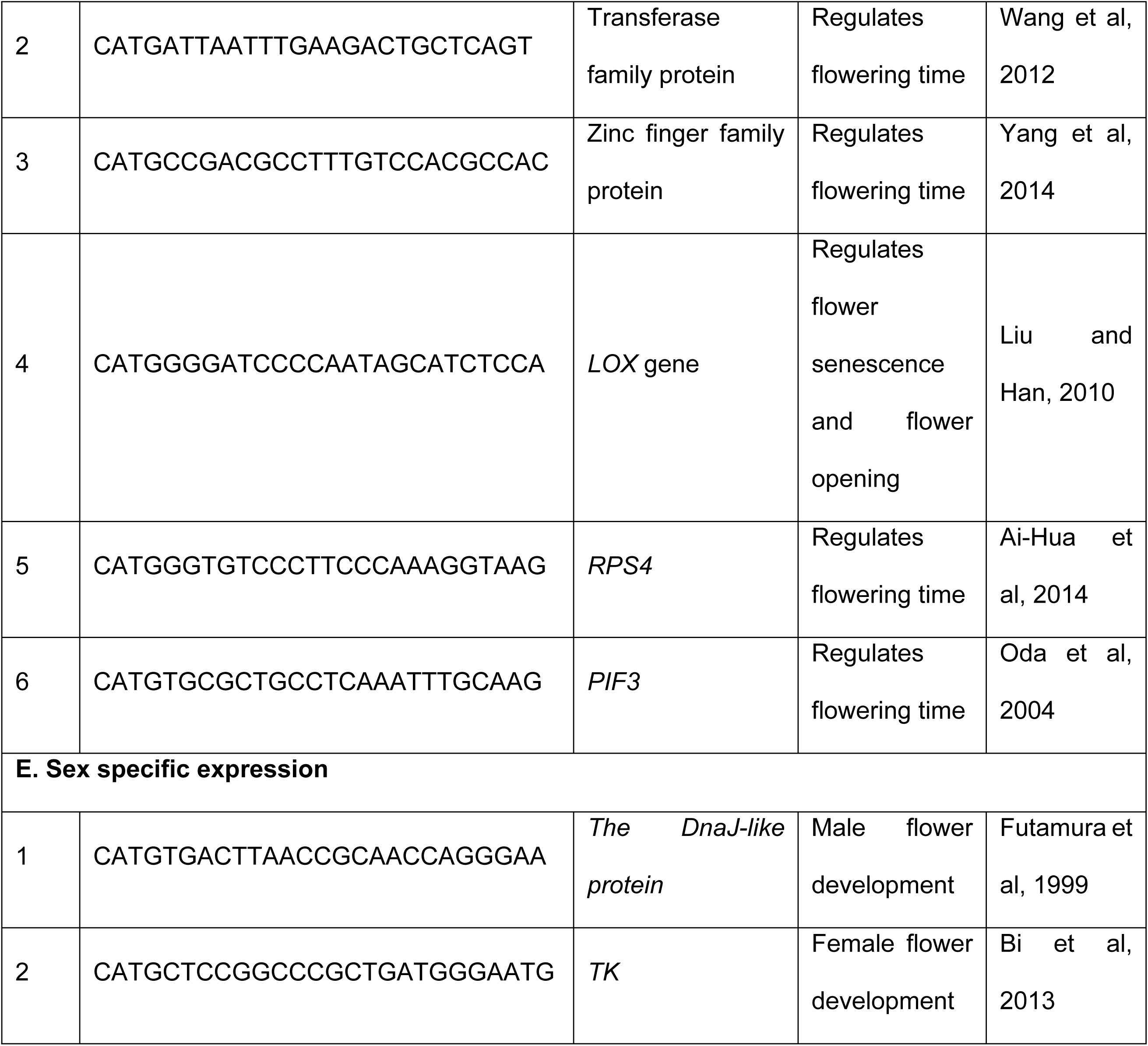
Summary of candidate genes with respective tag sequence and biological roles in flower development

In tomato, the long chain fatty acid CoA synthetase gene that is highly expressed in anther and petals, specifically in the sites subject to epidermal fusion [33], has been shown to be important for flower development, as lack of this gene impairs fertility and floral morphology. Similarly, the *GDSL esterase/lipase and APG* genes have been suggested for its potential involvement in flowering [41]. Among genes with a role in flower color development, our SuperSAGE analysis identified a *glutathione S-transferase-like* gene, reported for flower color intensity in *Dianthus caryophyllus* L [42].

Additional genes identified include those involved in flowering time, photoperiod and senescence regulation, such as: (1) a gene encoding NAC domain containing protein important for the coordination of cold response and flowering time [43], (2) a gene encoding a Transferase family *protein*, which is implicated in regulation of flowering time via the flowering repressor FLOWERING LOCUS C in *Arabidopsis thaliana* [44], a Zinc finger family *protein* gene involved in regulating flowering time and abiotic stress tolerance in *Chrysanthemum morifolium* [45], and a *LOX* gene known to regulate cell death related to flower senescence and flower opening [46,47]. A dramatic increase in *LOX* gene in response to senescence was observed in *Rosa hybrida* cv. Kardinal [40], in addition the *RPS4* gene is considered to play an important role in regulating flowering time, where a delay in flowering time was showed by silencing of the genes encoding *RPS4* and *Rhodanese* in *Glycine max* [48]. The transcription factor *PIF3* is considered to play an important role in the control of flowering through clock-independent regulation of CO and FT gene expression and is associated with early flowering in *Arabidopsis thaliana* [49]. Furthermore, genes known for organ specific expression were also identified including The DnaJ-like *protein* gene, detected predominantly in male flower of *Salix bakko* [50] and a Transketolase (TK) gene whose over-expression results in a higher ratio of female flowers and yield in cucumber [51].

In this study we have identified genes that are differentially expressed in early stage *D. rotundata* male, female and monoecious flowers, including genes previously reported as having roles in flowering and flower development. However, we recognize that the function (including in relation to sex determination) of these candidate flowering genes in yam is not known, and will require a combination of cloning and association studies in concert with functional studies (i.e. analysis of allelic variants such as null and hypomorphic alleles, and over and ectopic expression analyses). With the whole genome sequencing of *D. rotundata* recently released and the recent report of a protocol for genetic transformation in yam [52], such studies will provide powerful tools for genetic improvement of yam and dissection of the molecular networks controlling sex determination and flowering in yams.

### Validation of selected candidate genes associated with flowering in yams

In this study, we have identified genes that are differentially expressed, including genes previously reported as having roles in flowering and flower development, in early stage *D. rotundata* male and female flowers. Our validation tests of gene expression by qRT-PCR were in accordance with the SuperSAGE data for three of the four genes selected across samples representing male and female flowers. However, the result for DnaJ like protein showed discrepancy where the expression was higher in female than male flower but vice-versa for differential expression based on the next-generation sequencing based SuperSAGE technique. This discrepancy could be due to the samples used being not identical across both experiments.

## Conclusions

In conclusion, the current study is the first to identify candidate flowering-related genes in yam using a whole genome transcriptome profiling approach. Once verified with further experiments, the candidate sex determination and flowering associated genes identified in this study can be targeted for manipulation of flowering. Such genes provide important inputs for studies aiming to overcome the erratic to non-flowering nature of the yam crop, and thereby could contribute to improving breeding efficiency in yams. This is important in view of our observation that most *D. rotundata* accessions are either non-flowering or flowering is inconsistent from year to year. In addition to the need for further genetic and genomic studies on flowering and sex determination in yam, understanding the environmental and epigenetic factors controlling sex and flowering regulation in yam will also be important to underpin the improvement of breeding systems for yam.

## Acknowledgements

This work was conducted as part of the international collaborative research project “Use of genomic information and molecular tools for yam germplasm utilization and improvement for West Africa (EDITS-Yam)” funded by the Japan International Research Center for Agricultural Sciences (JIRCAS). GG acknowledges support from the Netherlands Ministry of Foreign Affairs, and a PhD International Merit Scholarship fee waiver from the National University of Ireland Galway.

## Supporting information

**S1 Table.** Flower related and other phenotypic traits used for characterization of *D. rotundata* accessions.

**S2 Table.** List of primers and their sequences used for qRT-PCR analysis.

**S3 Table.** List of all unique tags extracted from SuperSAGE library and used for analysis of differential expression.

**S4a Table.** List of differentially expressed tags between male and female flower group (p and FDR values< 0.01).

**S4b Table.** List of differentially expressed tags between male and monoecious flower group (p and FDR values < 0.01).

**S4c Table.** List of differentially expressed tags between female and monoecious flower group (p and FDR values < 0.01).

*logFC=Fold change of UP and DOWN expression; logCPM= log2(counts per million)/tag abundance; FDR=false discovery rate.

**S5 Table.** List of differentially expressed tags, sequence BLAST result and expression or involvement of the genes in flower and flower development.

**S6 Table.** Distribution of the differentially expressed tags across *D. rotundata* pseudo chromosomes.

**S7 Table.** Mean monthly temperature and rainfall during 2010 and 2011 yam growing seasons.

